# The polyQ expansion modulates the configuration and phosphorylation of huntingtin

**DOI:** 10.1101/721191

**Authors:** Taeyang Jung, Baehyun Shin, Giorgio Tamo, Hyeongju Kim, Ravi Vijayvargia, Alexander Leitner, Maria Jose Marcaida, Juan Astorga-Wells, Roy Jung, Ruedi Aebersold, Matteo Dal Peraro, Hans Hebert, Ihn Sik Seong, Ji-Joon Song

**Affiliations:** Department of Biological Sciences, Korea Advanced Institute of Science and Technology (KAIST), Daejeon 34141, Korea; School of Engineering Sciences in Chemistry, Biotechnology and Health, Department of Biomedical Engineering and Health Systems, KTH Royal Institute of Technology, S-141 52 Huddinge, Sweden; Department of Biosciences and Nutrition, Karolinska Institutet, S-141 83 Huddinge, Sweden; Center for Genomic Medicine, Massachusetts General Hospital, Boston, MA 02114, USA; Department of Neurology, Harvard Medical School, Boston, MA 02114, USA; Institute of Bioengineering, School of Life Sciences, Ecole Polytechnique Fédérale de Lausanne (EPFL), 1015 Lausanne, Switzerland; Department of Biology, Institute of Molecular Systems Biology, ETH Zurich, 8093 Zurich, Switzerland; Department of Medical Biochemistry & Biophysics, Karolinska Institutet 171 65 Solna, SWEDEN; HDxperts AB, 18348 Täby, Sweden; Faculty of Science, University of Zurich, Switzerland

## Abstract

The polyQ-expansion at the N-terminus of huntingtin (HTT) is the prime cause of Huntington’s disease. The recent cryo-EM structure of HTT with HAP40 provides information on the protein’s prominent HEAT-repeats. Here, we present analyses of the impact of polyQ-length on the conformation of HTT by cryo-EM, the domain-interactions by cross-linking mass spectrometry and the phosphorylation of HTT. The cryo-EM analysis of normal (Q23-) and disease (Q78-) type HTTs in their apo forms shows that the structures of apo HTTs significantly differ from the structure of HTT-HAP40, and that the polyQ expansion induces global structural changes consisting of significant domain movements of the C-HEAT domain relative to the N-HEAT domain. In addition, we show that the polyQ-expansion alters the phosphorylation pattern across the full-length HTT and that the specific phosphorylation (Ser2116p) in turn affects the global structure of HTT, which influences the activity of polyQ-expanded HTT. These results provide a molecular basis for the effect of the N-terminal polyQ segment on HTT structure and activity, that may be important for the cell-selective toxicity of mutant HTT.

## INTRODUCTION

Huntington’s disease (HD) is a progressive autosomal dominant neurodegenerative disease that is diagnosed by an uncontrolled motor movement known as chorea, cognitive disorder, and depression (Margolis and Ross, 2003; Walker, 2007). HD is caused by a CAG triplet repeat expansion, leading to an abnormal polyglutamine (polyQ) expansion in huntingtin (HTT) (Myers, 2004), but the mechanism of the disease development including how the expansion of only 30-40 glutamine repeat alters the function of HTT invoking HD, is still not understood. Multiple studies have revealed a proportional relationship between the length of the expanded repeat and the severity of disease (Lee et al., 2012; McNeil et al., 1997; Rubinsztein et al., 1996; Walker, 2007), indicating that the mechanism triggering the disease process involves a novel gain of function that is conferred on mutant HTT by the expanded polyQ tract (Gusella and MacDonald, 2000; Nucifora et al., 2001). HTT is expressed in most tissues with various expression levels and involved in a number of cellular functions (Sapp et al., 1997; Walker, 2007). Furthermore, the HTT gene is essential for embryonic development (Jeong et al., 2006). Therefore, it is critical to fully understand the effect of the polyQ region on HTT structure and function, including the normal function of HTT and the augmented toxic activity conferred on HTT, which will contribute to reveal the precise disease mechanism by which mutant HTT affects the highly complex HD pathogenic cascade.

The HEAT (<u>H</u>untingtin, <u>E</u>longation factor 3, protein phosphatase 2<u>A</u>, <u>T</u>arget of rapamycin 1) repeat motif is the major feature of HTT structure (Andrade and Bork, 1995). The structural flexibility of HTT as a HEAT repeat protein has been suggested in previous structural and biophysical studies (Grinthal et al., 2010; Halder et al., 2015; Yoshimura and Hirano, 2016) and is consistent with HTT’s interaction with various partners and participation in multiple cellular functions (Harjes and Wanker, 2003). This flexibility of HTT may be an important structural feature which is critical to its function and the polyQ tract could affect the overall structure rather than a limited region near the N-terminus. Indeed, we previously performed singleparticle EM analysis and analyzed intramolecular interactions of normal and mutant HTT with a panel of highly purified human recombinant HTTs, suggesting polyQ length dependent overall structural changes (Vijayvargia et al., 2016). Moreover, these structural differences of purified HTT caused by the length of the polyQ tract resulted in the enhancement of Polycomb Repressive Complex 2 (PRC2) histone methyltransferase activity in a cell-free assay (Seong et al., 2010). However, it needs to be further elucidated the mechanism by which the polyQ length expansion alters the spherical α-helical solenoid structure of HTT resulting in abnormal activity. Recently, a high-resolution cryo-EM structure of the HTT bound to HAP40 (HTT-associated protein 40) shows that HTT is composed of the N-HEAT, Bridge and C-HEAT domains, strongly supporting that HTT is a functional multivalent scaffold hub (Guo et al., 2018). Although this structure revealed the architecture of HTT consisting of three domains at the atomic level, it is not sufficient to understand how the flexible structure of HTT enables its multiple function without knowing the apo structure. Moreover, it is still difficult to determine whether the polyQ tract expansion may modulate the function because about 20% of the unresolved sequences of HAP40 bound HTT structure include the polyQ tract region and most of the posttranslational modification sites (PTM) both of which may be closely related to the flexible structure and very important for HTT function.

Here, we investigated the structural differences among HTTs in different states: HTT bound to HAP40, Q23-HTT and Q78-HTT, revealing that HTT has a dynamic but modular character and undergoes large conformational changes. Furthermore, we provide a proof-of-concept that polyQ expansion can induce HTT gain-of-function via interplay between phosphorylation status and conformational rearrangement of the domains of HTT.

## RESULTS

### Cryo-EM analysis on full-length Q23-HTT in the apo state

HTT is a modular protein containing HEAT repeats and the expansion of CAG repeats in the *HTT* gene causing polyQ track expansion in the HTT protein is a direct cause of HD. However, it has neither been shown polyQ expansion alters the global structure of HTT, nor if its conformation is affected by binding partners.

The recent atomic resolution structure (PDB: 6EZ8) showed that HTT is composed of three domains: N-HEAT, Bridge and C-HEAT with the Bridge domain connecting the two HEAT domains (Guo et al., 2018). HAP40 is bound between the N-HEAT and C-HEAT domains and seems to compact HTT, suggesting that HAP-40 binding may induce a conformational change of HTT. This observation is consistent with the idea that HTT is composed of flexible modular domains whose movement may be influenced by the polyQ expansion.

To investigate the structure of HTT in the apo form and its structural change, if any, upon the polyQ expansion, we first determined a cryo-EM structures of Q23-HTT. Monomeric Q23-HTT was isolated by 10-30% sucrose density gradient fractionation in the presence of disuccinimidyl suberate (DSS) crosslinkers (Figure S1). We collected 2,331 micrographs with a Titan Krios 300 keV electron microscope equipped with a Gatan Summit K2 direct detector using a Volta Phase Plate (VPP) (Figure 1A, Table S1). A total 72,065 particles from 790 selected micrographs were picked and the final 20,825 particles were used for 3D reconstruction using Relion2.1 (Figure 1B, 1C and S2). As estimated by gold-standard FSC, the final structure was obtained at 9.6 Å resolution (Figure S2C). The innate flexibility of HTT probably precluded reaching a higher resolution. To further validate the cryo-EM structure, we independently processed the identical particle set using cisTEM with an independent initial model generated from cisTEM (Grant et al., 2018). The cross-correlation coefficient (CCC) between the cryo-EM maps generated from Relion2.1 and cisTEM was 0.94 indicating that our cryo-EM map of Q23-HTT is likely to be free from processing artifacts (Figure S3).

**Figure 1.**
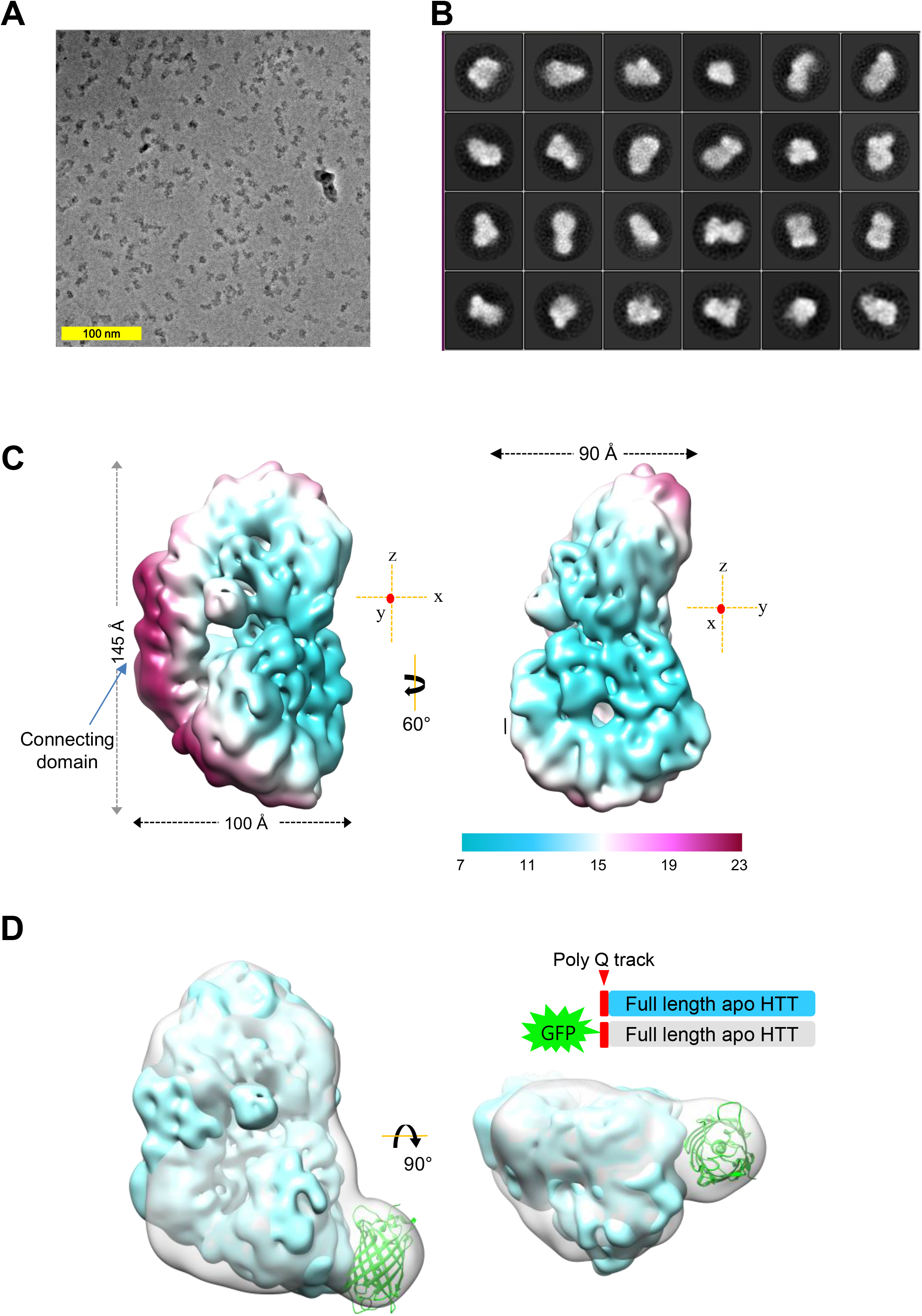
Cryo-EM analysis of full-length Q23-HTT in apo state. (A) A representative micrograph of Q23-HTT at a defocus of 0.7 μm recorded using a Volta phase plate. (B) 2D class averages of Q23-HTT particles used for 3-D reconstruction. (C) Cryo-EM map of Q23-HTT with color coding by locally estimated resolution in two different view angles. The dimensions of the Q23-HTT are 100 Å x 90 Å x 145 Å along the x, y and z axes respectively. A three-axis plot indicates the orientation (red dots indicate an axis perpendicular to the plane). (D) Superimposition of cryo-EM maps of Q23-HTT and GFP-Q23-HTT. GFP (PDB code: 1GFL) was fit in the extra density protruding from Q23-HTT indicating the position of N-terminus of Q23-HTT.

The refined Q23-HTT map shows that the overall dimensions of Q23-HTT are 145 x 90 x 100 Å, consistent with the previous structure obtained by negative-stain single-particle EM (Vijayvargia et al., 2016) (Figure 1C). The structure shows that Q23-HTT could accommodate the three domains: N-HEAT, Bridge and C-HEAT domains as defined from the high-resolution structure of HTT bound to HAP40. Overall, the global architecture of Q23-HTT is similar to that of HTT in complex with HAP40.

As the cryo-EM map at the current resolution did not enable us to independently locate the N-terminus of Q23-HTT where the polyQ expansion occurs, we aimed to map the N-terminus by determining the cryo-EM structure of N-terminal GFP-tagged HTT (GFP-HTT). GFP is a compact protein and has been utilized for mapping the locations of domains in large complexes in low resolution cryo-EM maps (Ciferri et al., 2015). We generated GFP-HTT and determined its cryo-EM structure. The final refined 3D map of GFP-HTT was reconstructed from 17,689 particles with an overall 15.3 Å resolution. Superimposition of GFP-HTT on Q23-HTT (CCC: 0.88) revealed an apperent extra density (Figure 1D, Figure S4), clearly indicating the location of the N-terminus of Q23-HTT.

To gain further insight into the structure of Q23-HTT in the apo state, we generated and fitted atomic models to our cryo-EM map of Q23-HTT utilizing the structure of HTT in complex with HAP40 (HTT_HAP40_) via an integrative modeling strategy. First, the HTT_HAP40_ model was split into three rigid domains: N-HEAT, Bridge and C-HEAT. Then, the domains were rigidly fitted into the Q23-HTT density map using a hierarchical multi-body constrained molecular docking strategy (Tamo et al., 2017). First, the best fitting N-HEAT domain from the rigid fitting was selected, which was eventually fully consistent with the location of GFP in GFP-HTT. The bridge domain and C-HEAT domain were subsequently placed on the map via multi-body fitting. The final representative model generated in this first stage showed an overall good fit to the density map (CCC value of 0.71), which was further improved by a MD-based flexible refinement using Molecular Dynamics Flexible Fitting (MDFF) (Figure S5A). The final MDFF-refined Q23-HTT model has a CCC of 0.86 with the cryo-EM map (Figure 2A). The Q23-HTT atomic model contains three distinctive domains, representing HEAT repeat motifs with a reliable fitting score to the experimental cryo-EM map (Figure 2B).

**Figure 2.**
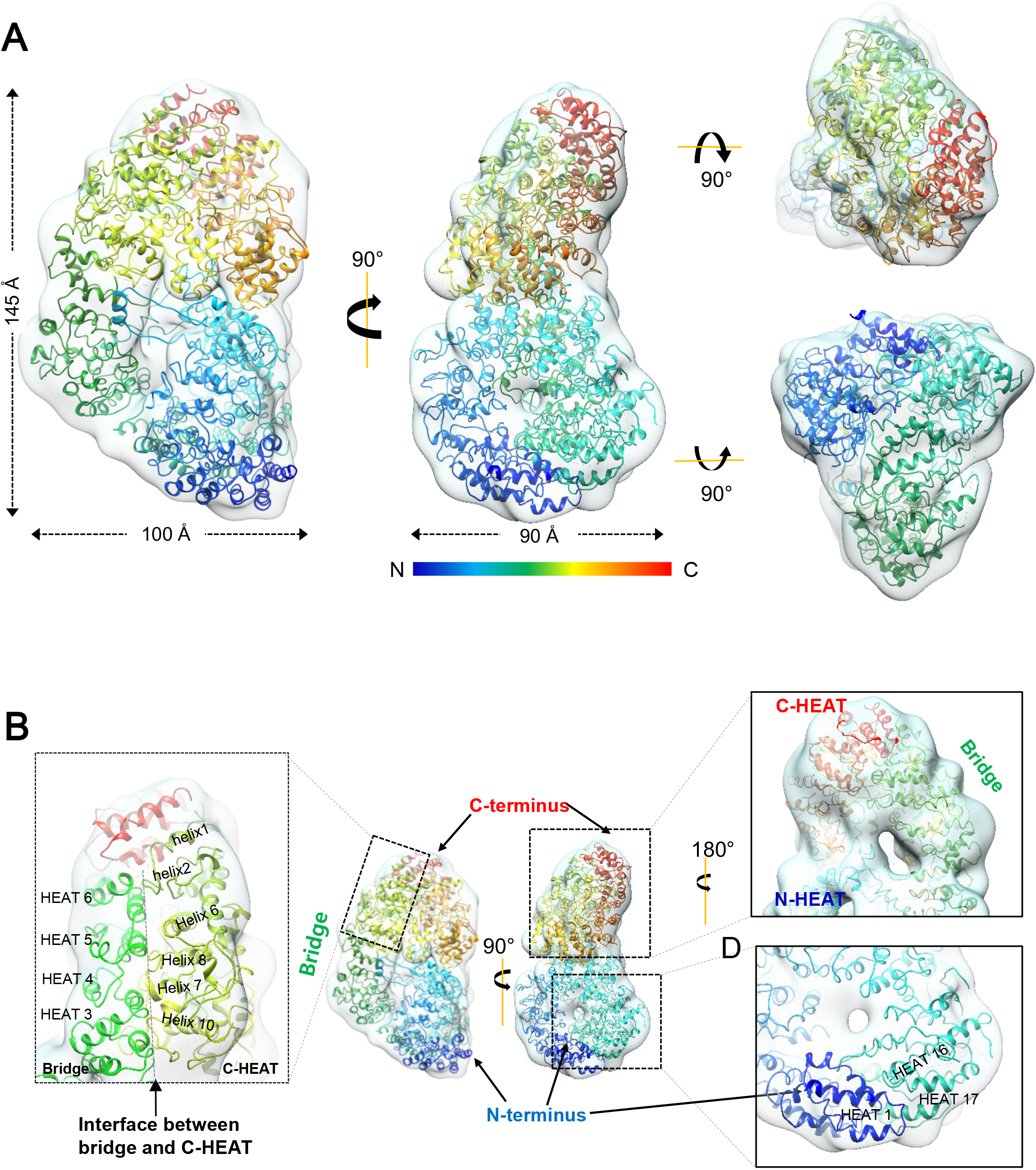
Modeling of the atomic structure of Q23-HTT based on the HTT-HAP40 complex. (A) Q23-HTT MDFF model superimposed on the cryo-EM map of Q23-HTT. The Q23-HTT model is shown rainbow-colored from N- to C-terminus in a cartoon model. (B) Detailed view of the Q23-HTT model fitted on the cryo-EM map. The N- and C-HEAT domains are colored in rainbow, and the BRIDGE domain in green.

### Conformational Change of the C-HEAT domain in HTT

Having an Q23-HTT model based on our cryo-EM map, we compared Q23-HTT and HTT_HAP40_ to examine their structural differences (Figure 3A). The overall dimensions significantly differ between the two structures. Q23-HTT is more extended along its longest axis (the z-axis defined in Figure 3B), while it is rather compact in the presence of HAP40 (Figure 3A). To compare the two structures in details, we aligned Q23-HTT and HTT_HAP40_ by superimposing the N-HEAT domains (Figure 3B and 3D). This practice revealed a significant difference in the position of the C-HEAT domain relative to the N-HEAT domain, which grabs HAP40 with the N-HEAT domain (Figure 3B and 3C). To evaluate the relative movement among the three domains in greater detail, we first measured the movement of the Bridge domains by anchoring the N-HEAT domains. With the beginning of the Bridge domain serves as a pivot joint, the Bridge domain of Q23-HTT is rotated about 20° along the y-axis (perpendicular to the plane as viewed in Figure 3B) and 60° along the z-axis, compared to the Bridge domain of HTT_HAP40_ (Figure 3B Left panel). Next, we measured the movement of the C-HEAT domain by superimposing the Bridge domains (Figure 3B Right panel). The end of the Bridge domain serves as another pivot point and the C-HEAT domain of Q23-HTT is rotated about 50° and rolled over about 35°, compared with the C-HEAT domain of HTT_HAP40_ (Figure 3C). When these two movements are combined, the C-HEAT domain is rotated 20° at the y-axis, 110° at the z-axis and 35° rolled over (Figure 3C). There are two pivot points in HTT at the beginning and end of the Bridge domain as if HTT is a modular structure. With these movements, the CHEAT domain is consequently moved away from the N-HEAT in Q23-HTT compared with the HTT_HAP40_. This structural comparison indicates that the positional change of the C-HEAT domain relative to the N-HEAT domain may have an impact on the function of HTT by modulating its interactions with HAP40. While the relative orientations of the three domains of Q23-HTT and HTT_HAP40_ differ significantly, the conformation within each domain is similar with subtle difference (Figure 3D). In short, HTT exhibits a large C-HEAT domain swing-out movement by utilizing two pivot points at the beginning and end of the Bridge domain, when the N-HEAT domains are aligned. These data show that HTT has a flexible and modular character, and suggest that the conformational change among the domains may play a critical role on the function of HTT such as interacting with other proteins.

**Figure 3.**
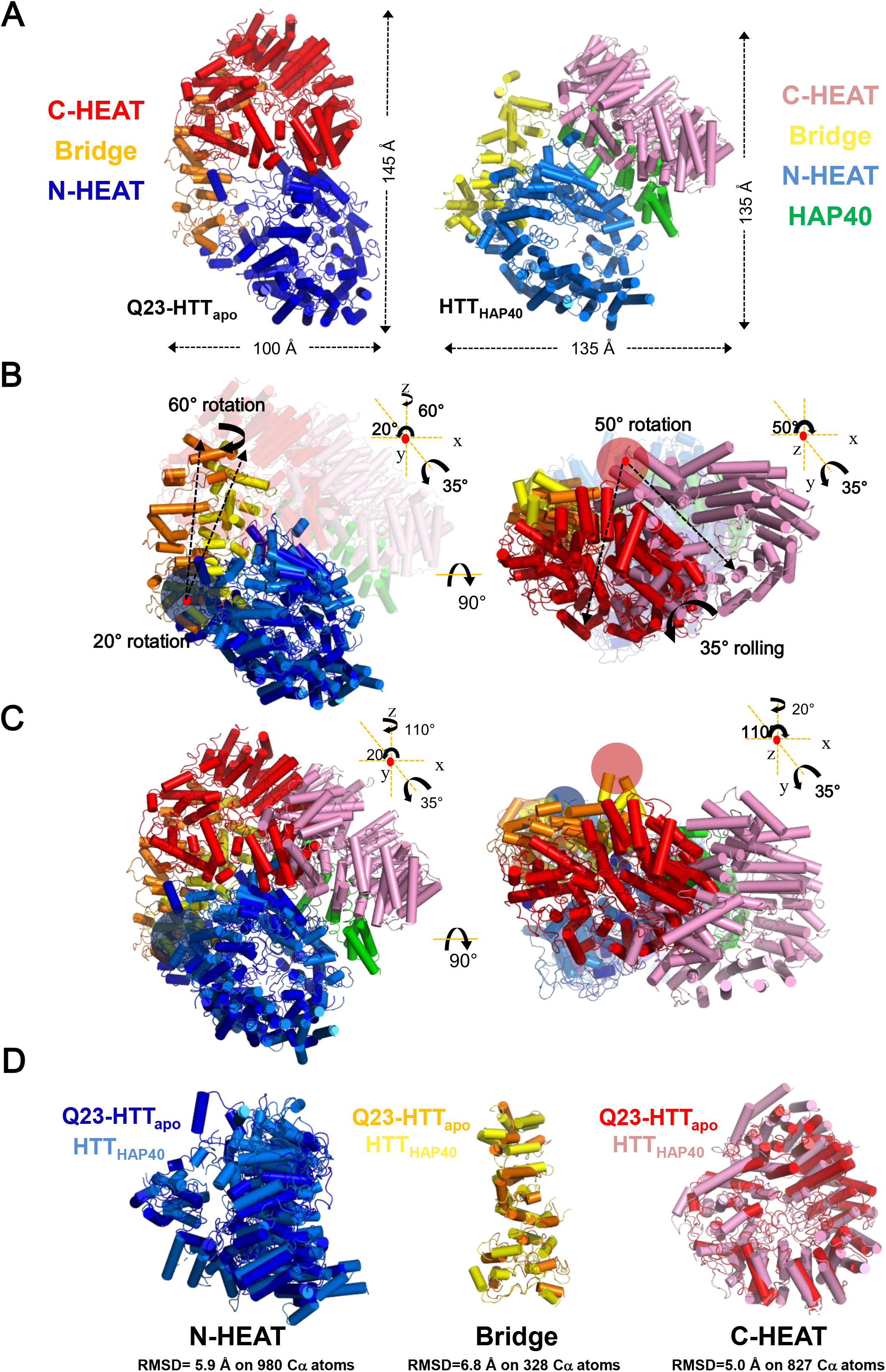
Conformational change of the C-HEAT domain in Q23-HTT compared with HTT_HAP40_. (A) Overall structures of Q23-HTT and HTT_HAP40_. N-HEAT, C-HEAT and Bridge domains are colored in blue/royal blue, orange/yellow and red/pink respectively. (B) N-HEAT domains of Q23-HTT and HTT_HAP40_ were superimposed revealing the difference in the relative positions of the Bridge domains (Left panel). The Bridge domain of Q23-HTT is rotated about 20° at y-axis (red dot, perpendicular to the plane), and rotated 60° along the z-axis, compared with HTT_HAP40_ (Left panel). The Bridge domains were superimposed revealing the difference in the relative positions of the C-HEAT domains (Right panel). The C-HEAT domains of Q23-HTT is rotated about 50° at z-axis, and rolled over about 35°. A three-axis plot with angles shows the movement of the domains. Dark blue and pale red circles indicate pivot joints. (C) Overall conformational change of Q23-HTT and HTT_HAP40_ structure. A three-axis plot with angles shows the movement of the domains at the pivot joints marked with dark blue and pale red circles. (D) Superimposition of three domains of Q23-HTT (cyan, orange and red for N-HEAT, Bridge and C-HEAT domain) on the corresponding domains of HTTHAP40 (royal blue, yellow and pink for N-HEAT, Bridge and C-HEAT domain). The root mean square distance (RMSD) values between each domain of Q23-HTT and their counter parts in HTT_HAP40_ were calculated and are shown in below.

### Structural comparison between Q23-HTT and Q78-HTT

The polyQ expansion at the N-terminus of HTT leads to HD, and accumulating data suggest that the polyQ expansion alters the global structure of HTT (Cui et al., 2014). We previously observed the structural differences between Q23-HTT and Q78-HTT using cross-linking mass spectrometry (XL-MS) and negative-stain EM (Vijayvargia et al., 2016). However, the precise nature of the structural changes caused by the polyQ expansion remains unknown. To gain detailed insights into the structural differences between Q23- and Q78-HTTs, we aimed to determine the cryo-EM structure of Q78-HTT. Q78-HTT protein was prepared in the same way as Q23-HTT and 1,552 micrographs were collected using a Titan Krios 300 KeV microscope with a Gatan Summit K2 detector using VPP (Figure 4A). A final set of 21,983 particles from the total of 45,038 particles picked were selected for subsequent 2D class-averaging and 3D reconstruction, generating a cryo-EM map of Q78-HTT at a resolution of about 20 Å (Figure 4A, 4B and S6). The overall dimensions of Q78-HTT are similar to those of Q23-HTT, featuring only a slight expansion alone the z-axis. Despite the relatively low resolution, we were able to clearly locate the position of the N-HEAT and C-HEAT domains. To compare the structure of Q78-HTT to that of Q23-HTT, we fitted the atomic model of each N- and C-HEAT domains onto the Q78-HTT cryo-EM map and refined them using MDFF. While the N- and CHEAT domains could be positioned within the map, we were not able to place the Bridge domain by rigid body fitting without deforming its structure, suggesting that there may be some substantial structural changes in the Bridge domain. Therefore, we fitted only the N- and CHEAT domains into our cryo-EM map. We then compared the relative orientations of N- and CHEAT domains of Q23- and Q78-HTTs. When two maps of Q23- and Q78-HTT were superimposed, there was a clear difference in the position of the C-HEAT domains relative to the N-HEAT domains. Specifically, the C-HEAT domains are rotated about 55° along the y-axis and rotated about 30° alone the z-axis with 35° rolling, with respect to each other (Figure 4C). The Bridge domain seems to serve as pivot joints as observed between Q23-HTT and HTT_HAT40_. These observations suggest that the polyQ expansion in the N-terminus of HTT may induce a change in orientation of the C-HEAT domain relative to the position of the N-HEAT domain.

**Figure 4.**
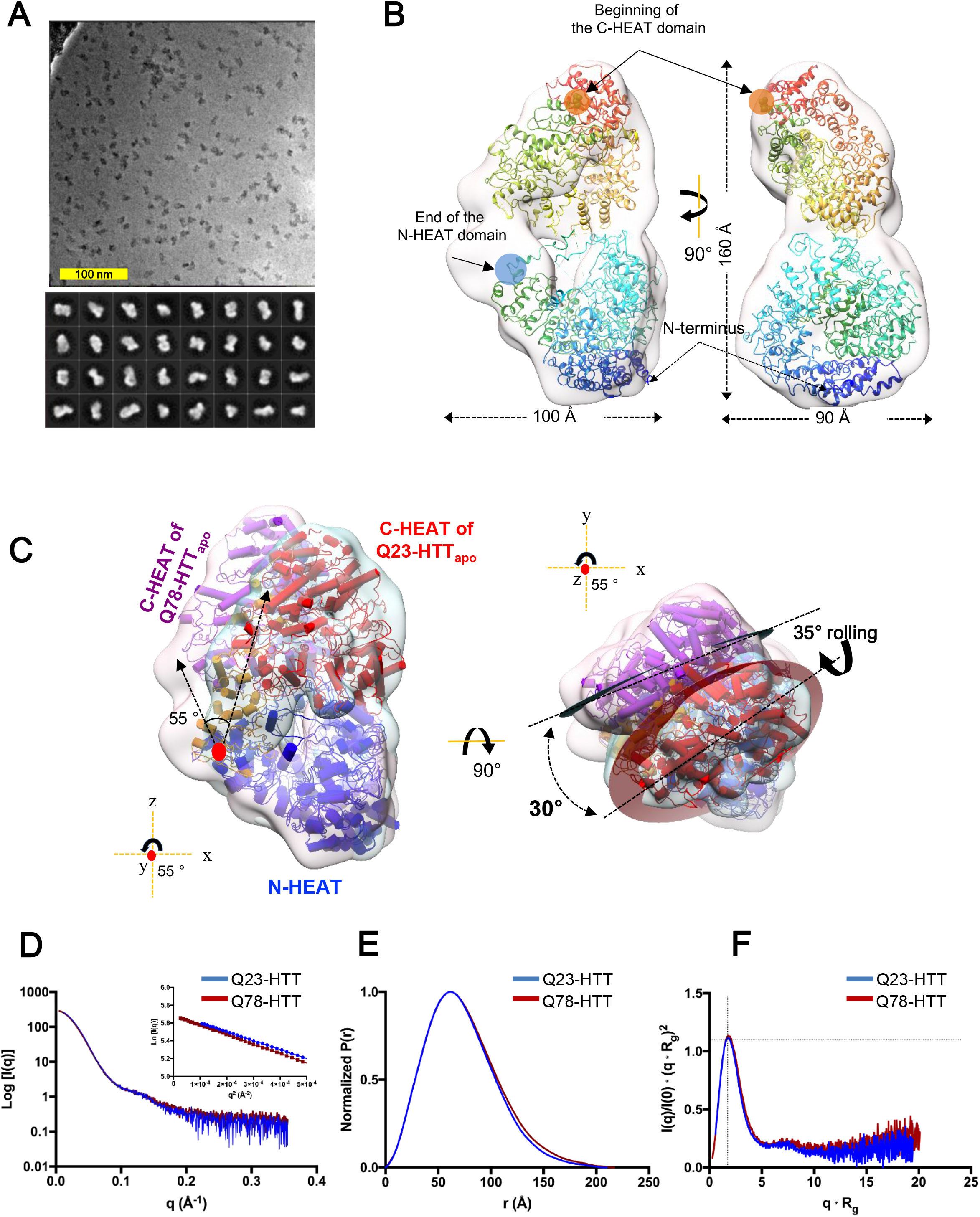
Cryo-EM analysis of full-length Q78 HTT in apo state and comparison with Q23-HTT in apo state. (A) A representative micrograph of Q78-HTT at a defocus of 0.5 μm recorded using a Volta phase plate (Top). 2D class averages of selected Q78-HTT particles used for single particle reconstruction (Bottom). (B) Cryo-EM map of Q78-HTT fitted with N- and C-HEAT domains by MDFF. The model is depicted in rainbow colors and fit-in the cryo-EM map. The cryo-EM map of Q78-HTT shows overall dimensions as 100 x 90 x 160 Å. The end of the N-HEAT domain and the pivot joints at the beginning of N-HEAT and the end of C-HEAT are indicated by blue and red circles respectively. (C) A superimposition of cryo-EM maps and models of Q23-HTT and Q78-HTT. N-HEAT domains of Q23-HTT and Q78-HTT are aligned revealing structural difference between the two structures. N-HEAT is shown in blue, Bridge in orange and C-HEAT domains are in red for Q23-HTT and magenta for Q78-HTT. Red dots show a y-axis (Left panel) or z-axis (Right panel), which are perpendicular to the surface. (D) One-dimensional intensity plot with the Guinier fit as insert. (E) Normalized Pair-distance distribution function, used to determine the maximum dimension of the particles (D_max_). (F) Dimensionless Kratky Plot which quantifies the conformational flexibility of proteins. The dotted lines are drawn at qRg = 1.73 and I(q)/I(0) ▪ (qRg)^2^=1.1. Folded proteins have a local maximum where the two lines intersect.

To further investigate the overall structural conformation and possible differences between Q23-, and Q78-HTT in solution, we performed Small Angle X-ray Scattering (SAXS) analysis. Scattering curves were obtained from the samples as they were eluting from the size exclusion chromatography column to avoid contamination from possible higher order oligomers (Figure 4D). This allowed for a good determination of the Guinier derived radius of gyration (*R*_g_) values for the Q23-, and Q78-HTT, which are 54.9 and 56.6 Å, respectively. The comparatively compact shape of the P(r) distribution functions (Figure 4E) indicates that the proteins are globular in solution, in agreement with the type determined by the software *DATCLASS* (Franke et al., 2017) (Table S2), which was “compact”, as well as cryo-EM maps. In addition, the dimensionless Kratky plot shows a maximum at the expected position for globular proteins (Figure 4F). The maximum dimension of the particles estimated from the P(r) distribution functions (D_max_= 210 Å for Q23-HTT and 217 Å for Q78-HTT) indicate that the proteins behave as monomers in solution and that Q78-HTT is slightly more extended than Q23-HTT, which is consistent with the models obtained through cryo-EM map fitting. The variability in the extension of the polyQ region between the two constructs is reflected in the differences in the R_g_ and D_max_ values observed (Table S2), which is also consistent with our cryo-EM analysis on Q23- and Q78-HTT in that Q78-HTT adopts a slightly more extended conformation compared with Q23-HTT. In short, the SAXS data consistently represented a globally similar ‘compact’ conformation of Q23- and Q78-HTT, as well as a slightly more extended dimension of Q78-HTT at the same time. Our comparative structural analysis between Q23- and Q78-HTTs suggests that there is significant conformational change in the relative orientations of the three domains, but not within the N- and C-HEAT domains. To further assess the conformational dynamics of Q23-, and Q78-HTT, we performed hydrogen-deuterium exchange mass spectrometry (HDX-MS). Q23- and Q78-HTT were incubated for 10, 45, 90 and 180 min for deuteration. The samples were then subject to protease digestion followed by mass spectrometry. We only analyzed peptides found in both Q23- and Q78-HTT and the peptide coverage for Q23- and Q78-HTT was 42.7% in both cases (Figure S7A and S7B). The deuterium exchange rate was plotted in a butterfly plot (Figure S8A). Despite the conformational change between Q23- and Q78-HTT observed in the cryo-EM analysis, there was no significant differences in deuterium exchange rate between Q23-, and Q78-HTT, which is consistent with our cryo-EM structural analysis showing that HTT is a dynamic modular structure composed of three domains and that there is no substantial structural difference within the modules. We mapped the locations of the peptide obtained from HDX-MS on the primary sequences and the structures of Q23- and Q78-HTT (Figure S7 and S8B). Peptides showing high deuterium exchange rate are clustered at several regions. The large region around a.a. 400-600 located within the N-HEAT domain shows the highest exchange rate and is disordered in cryo-EM structures. Interestingly, many PTMs are found in this region suggesting that the highly solvent accessible region may serve as a functional site modulated by PTMs (Rangone et al., 2004; Ratovitski et al., 2017; Schilling et al., 2006; Warby et al., 2005). Another region showing high exchange rate is around a.a. 2,000-2,200 and is located at the interface between the Bridge and C-HEAT domains, which is consistent with our observation that the Bridge domain serves as pivot joints making HTT conformationally flexible. Overall these data show that HTT is a dynamic modular structure and that the polyQ expansion at the N-terminus of HTT modulates the relative orientation of the C-HEAT domain, which may be implicated in mutant HTT functions.

### The conformational flexibility of HTT

The orientation of the C-HEAT domains in Q23- and Q78-HTT relative to the N-HEAT domain significantly differs suggesting that polyQ expansion may induce this structural change. In addition, our structural analysis revealed conformational differences between Q23-HTT in the apo state and HTT in complex with HAP40. To compare the dynamics observed among Q23-HTT, Q78-HTT and HTT_HAP40_, we placed the three structures with the N-HEAT domains superimposed, revealing that there is a continuum in the movement of the C-HEAT domain among HTT_HAP40_, Q23-HTT and Q78-HTT structures (Figure 5A and 5B). C-HEAT in HTT_HAP40_ is most closely located to the N-HEAT, resulting in the most compact structure due to the binding of HAP40 binding. In contrast, Q78-HTT adopts the most extended conformation, in that the C-HEAT domain is far away from the N-HEAT domain. The C-HEAT domain in Q23-HTT is located in the middle of these two conformations (Figure 5A). This analysis indicates that HTT has two pivot joints located at the beginning and at the end of the Bridge domain, which leads to the dynamic nature of HTT. Specifically, at the joint at the beginning of the Bridge domain (blue circles in Figure 5B), the Bridge domain can rotate along the y-axis and can roll at the longest axis. At the joint at the end of Bridge domain (pale red circles in Figure 5B), the C-HEAT domain rotates along the y-axis and rolls. It is worth noting that the CHEAT domain is located in closer proximity to the N-HEAT domain domain than Q23-HTT, which is caused by HAP40 binding, and that the relative location of the C-HEAT domain to the N-HEAT domain is altered upon the polyQ expansion. This observation suggests that the changes in the relative orienations among thedomains may modulate HTT function.

**Figure 5.**
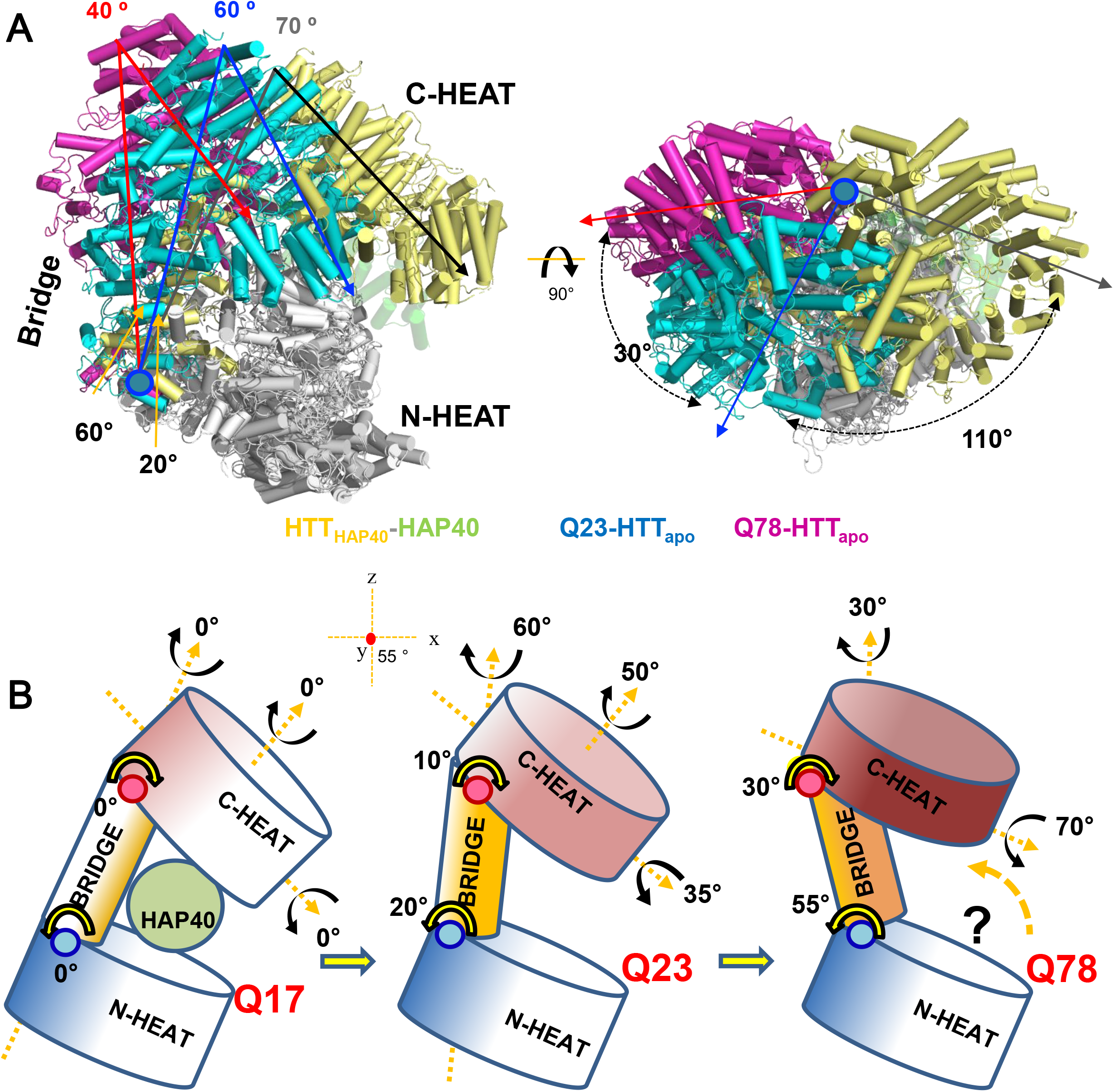
Conformational flexibility of the C-HEAT domain of HTT. (A) Superimposition of Q23-HTT, Q78-HTT and HTT_HAP40_ models aligning N-HEAT domains. Left panel: the Bridge and C-HEAT domains colored in cyan for Q23-HTT, pink for Q78-HTT and pale yellow for HTT_HAP40_. For clarity, the N-HEAT domains were colored in white. The rotation angles at the pivot joint (pale blue circle) located at the beginning of Bridge domain are indicated below, and the angles between the Bridge and C-HEAT are shown above. Right panel: 90° rotated view from the left panel. The rotation angles of C-HEAT at the pivot joint (pale blue circle) are indicated. (B) A schematic model for conformational flexibility of HTT. The angles in HTT_HAP40_ are set 0° to compare relative positions in Q23- and Q78-HTT. N-HEAT domains are shown in blue, Bridge in yellow and C-HEAT in red. Pale blue and pale red circles indicate two pivot joints.

### Polyglutamine tract length alters phosphorylation patterns along the entire protein

The 3,144 residue Q23-HTT is predicted to possess many potential phosphorylated serine (S), threonine (T) or tyrosine (Y) residues (Blom et al., 2004). In fact, more than 70 phosphorylation sites across the entire protein have been collectively reported from different studies with human cells (Hornbeck et al., 2012), implying that phosphorylation of HTT can be significantly involved in its function. Since the overall structure of HTT can be affected by the polyQ tract expansion (Figure 4) but many known phosphorylation sites were not determined in any of HTT cryo-EM structures, we decided to investigate how the phosphorylation patterns of HTT are affected by the polyQ expansion. To evaluate the phosphorylation status at sites consistently identified in full-length HTT, we have utilized Sf9 insect cells, an excellent surrogate for the mammalian system with many kinase families conserved (Busconi and Michel, 1995), to express recombinant full-length HTT with various polyQ lengths (Seong et al., 2010). To identify phosphorylated peptides, we purified Q23-, Q46- and Q78-HTT, in the presence of phosphatase inhibitors, and performed three independent tandem mass spectrometry (LC-MS/MS) analyses. Alignment of the HTT peptides revealed high coverage (~80% of 3,144 residues) and a generally similar set of peptides with phosphorylated serine or threonine residues (Figure 6A and S9A). Sixteen phosphorylation sites were identified more than twice.

**Figure 6.**
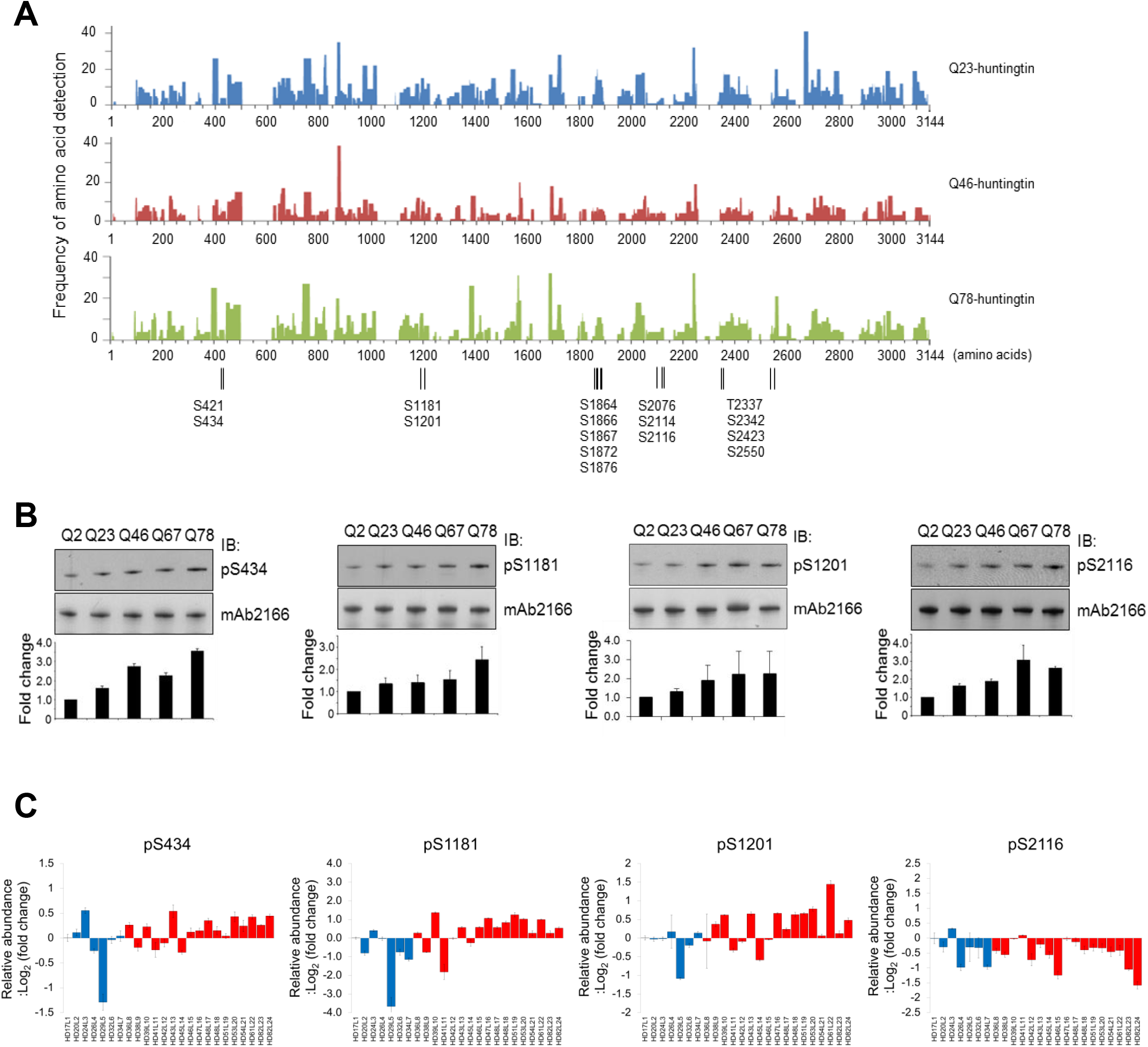
Polyglutamine length dependent alteration of phosphorylation in purified HTT protein and human cells. (A) Coverage graphs of full-length HTT proteins subjected to mass spectrometry (LC-MS/MS) analysis for determination of phosphorylation sites. Both normal (Q23-HTT) and mutant (Q46- and Q78-HTT) proteins show similar coverage patterns and to the same extent (~80%). Several HTT phosphorylation sites were identified by MS and are listed along the length of the HTT amino acid sequence. (B) The purified recombinant Q2-, Q23-, Q46-, Q67-, and Q78-HTT series were subjected to immunoblot analysis and probed with a panel of 15 phosphorylation site specific antibodies. The results of four phosphorylation sites (pS434, pS1181, pS1201 and pS2116) are shown here along with the quantification graph. The signal was normalized to total HTT levels (the signal of mAb2166). (C) A panel of 24 lymphoblastoid cell lines (7 controls, blue bar and 17 HD subjects, red bar) was subjected to PRM-based targeted mass spectrometry experiments in order to ascertain the profile of HTT phosphorylation in human cells. The three targeted ions derived from phosphopeptide containing pS434, pS1181, pS1201 and pS2116 were extracted and summed to represent the relative abundance of phosphopeptides (Y-axis, log2 scale). Data represent mean of three experiments (technical replicates).

They included 15 previously reported and one novel site (S2423) (Aiken et al., 2009; Anne et al., 2007; DeGuire et al., 2018; Dephoure et al., 2008; Dong et al., 2012; Hornbeck et al., 2012; Huang et al., 2015; Humbert et al., 2002; Luo et al., 2005; Moritz et al., 2010; Phanstiel et al., 2011; Rangone et al., 2004; Ratovitski et al., 2017; Schilling et al., 2006; Warby et al., 2005), confirming the utility of insect cells for studying phosphorylation patterns in HTT. Most phosphorylation sites except for S2114, S2116, S2423 and S2550 located in the C-HEAT domain are located in disordered regions in the cryo-EM structures.

To systematically examine the potential impact of polyQ tract length on the pattern of HTT phosphorylation, we first generated a panel of phosphosite-specific affinity purified antibodies for each of these 16 sites, demonstrating their site and phosphorylation specificities in a dot-blot assay using the phospho and the corresponding non-phospho peptides (Figure S9B). Using this panel of phosphosite-specific antibodies, immunoblot analysis of the purified recombinant Q2-, Q23-, Q46-, Q67-, and Q78-HTT series demonstrated that for 8 reagents, the band intensities (normalized to total HTT detected by mAb2166 and relative to Q2-HTT) increased with polyQ tract length (Figure 6B, S10, S11 and Table S4). This finding reveals that the size of the polyQ tract affected the level of HTT phosphorylation at sites (S421, S434, S1181, S1201, S1864, S1876, S2116 and T2337) across the entire protein instead of a cluster near the polyQ tract. It is plausible that the effect of polyQ expansion on the PTM is due to the global conformational change of full-length HTT potentially through altering the relative position of the C-HEAT domain (Figure 5).

We then investigated the phosphorylation at 16 sites in endogenous HTT from HD patient cells to understand their phosphorylation levels and patterns depending polyQ tract length in human cells. We chose lymphoblastoid cell lines (LCL) which most highly express *HTT* mRNA (GTEx portal). We could detect specific signals from LCL lysates by using our HTT phospho antibodies (data not shown) but the phospho antibody signals were too variable to obtain statistically significant patterns with a few LCL lines likely due to the dynamic nature of cellular phosphorylation. For accurate and reliable quantification in multiple lines at the same time, we utilized a parallel reaction monitoring (PRM) strategy, a highly sensitive and quantitative targeted MS/MS technique, to quantify the phosphorylation level of each site in cellular HTT. We found seven phosphopeptides suitable for the PRM assay (Figure S12). Six of these sites (pS421, pS434, pS1181, pS1201, pS1876 and pS2116) were significantly affected by polyQ tract length in the allelic series of purified HTT while pS1872 was not. Using the PRM assay, we successfully identified and quantified substantial amounts of these seven phosphopeptides in endogenous HTT from all 24 LCLs expressing HTT with different polyQ lengths (Figure 6C and S13), confirming phosphorylated HTT at these seven sites in human cells. In order to examine the impact of the polyQ tract length on these phosphosites, we performed polyQ length correlation linear regression analysis with their relative abundances and two sites, S1201 and S2116, showed nominal P-values (Table S4). The phosphorylation levels of S434 and S1181 also seemed to be associated with polyQ tract length, although their P-values were slightly higher than the nominal P-value. All four phosphorylation sites are significantly polyQ length dependent in purified HTT (Figure 6B and 6C), suggesting a common impact of polyQ expansion on phosphorylation, likely through changes in HTT’s structure. However, cellular context also appears to be a crucial determinant of HTT phosphorylation because the direction of pS2116 correlation with *HTT* CAG size in LCL was opposite in insect cells. Together, our results suggest that HTT phosphorylation and polyQ expansion are interrelated through structure and our panel of purified human HTT with different polyQ tract sizes, and provides basis to further study the effect of polyQ expansion on HTT structure and phosphorylation.

### S2114/S2116 phosphorylation in the C-HEAT domain related to HTT structure and activity

We then determined whether phosphorylation status might influence HTT structure and/or its functional activity. We previously showed that polyQ expanded HTT (Q78) stimulates PRC2 histone methyltransferase activity (Seong et al., 2010). Notably, this assay has previously been validated as a read-out for normal function of full-length HTT (and the effect of its polyQ tract) on PRC2-dependent chromatin marks and regulation as demonstrated both *in vivo* (Seong et al., 2010) and in cell culture studies (Biagioli et al., 2015; Shin et al., 2018). Using this assay system as a readout, we examined the effect of phosphorylation on HTT function. In order to narrow down specific phosphorylation sites that may affect the PRC2 enhancing activity of mutant HTT, we selected S434, S1181, S1201, S2114 and S2116 sites exhibiting increased levels of phosphorylation as the length of polyQ increases, and then purified five alanine mutants of both Q23- and Q78-HTT at S434, S1181, S1201, S2114 and S2116 (Figure S14). Two out of five alanine mutant Q78-HTTs, S2114A and S2116A exhibited significantly reduced activity compared to the Q78-HTT control (Figure 7A). Moreover, when we performed the PRC2 assay with Q23 S2116A-HTT and Q78 S2116A-HTT, the alanine mutation on S2116 dampened the extra PRC2 stimulating activity of Q78-HTT, whereas the same mutation in Q23-HTT apparently did not affect it (Figure 7B). Removing the phosphorylation at S2116 site, therefore, removes the “extra” activity of Q78-HTT without losing the basal activity seen in Q23-HTT. Considering that the activity gained by the polyQ expansion at the N-terminus was compromised by eliminating the phosphorylation of S2116 located within the C-HEAT, there may be a functional interaction between the N-HEAT and the C-HEAT of HTT, which may be critical for interaction with PRC2 and is affected by the S2116A mutation. To test whether the mutation disrupted the interaction between PRC2 and HTT, we performed immunoprecipitation assays of reconstituted HTT-PRC2 complex with EZH2, the PRC2 catalytic subunit. The S2116A mutation mildly increased the interaction between Q23-HTT and EZH2, whereas the same mutation in Q78-HTT dramatically decreased the interaction between Q78-HTT and EZH2, while control Q78-HTT robustly immunoprecipitated (Figure 7C). These results suggest that S2116 phosphorylation uncovers a novel property of mutant HTT related to the interaction with PRC2.

**Figure 7.**
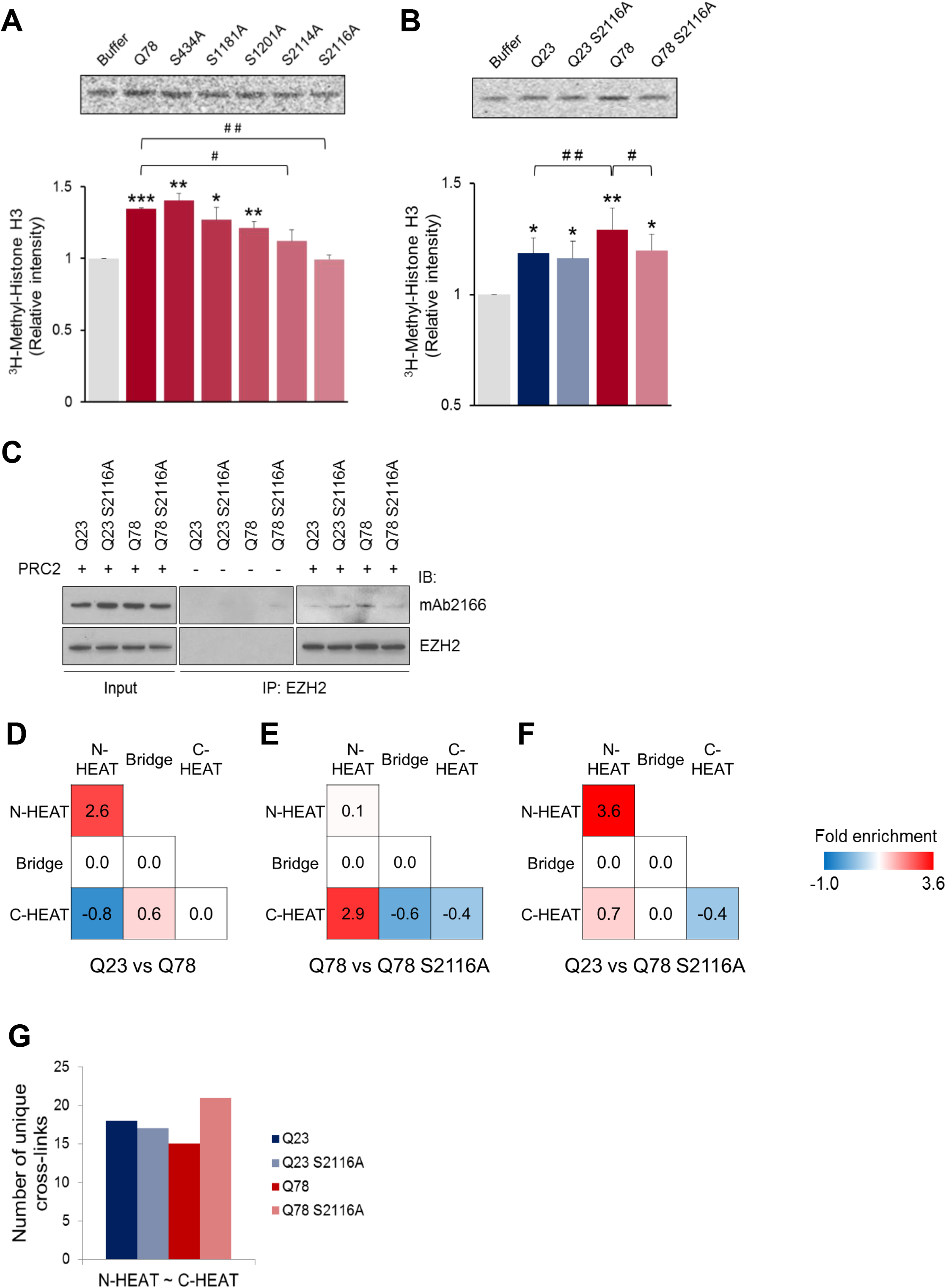
Phosphorylation status modulates HTT activity in a polyglutamine length dependent manner: the critical role of pS2116 on mutant HTT activity. (A) Cell-free PRC2 assay of five alanine mutants of Q78-HTT at S434, S1181, S1201, S2114 and S2116 showed that the enhanced activity of Q78-HTT was significantly decreased in both S2114A and S2116A mutants compared to wild-type Q78-HTT. (B) Cell-free PRC2 assay of two S2116A mutants of either Q23- or Q78-HTT revealed that the S2116A mutation affects only the extra activity of mutant HTT without altering a baseline activity. For both cell-free PRC2 assay (A and B), data are averages of three experiments and error bars are SEM. The significance was calculated and displayed compared to PRC2 alone (*, *P* < 0.05, **, *P* < 0.01, ***, *P* < 0.001), or as indicated (#, *P* < 0.05, ##, *P* < 0.01) using Student’s *t*-test. (C) Effect of HTT S2116 phosphorylation status on its interaction with PRC2 was analyzed by immunoprecipitation with anti-EZH2 antibody followed by immunoblotting for HTT. (D, E and F) Q23- and Q78-HTT including corresponding S2116 alanine mutants were cross-linked by DSS and intra-domain cross-linked peptides were identified by mass spectrometry. (D,E,F) The fold enrichment score between Q23- and Q78-HTT (D), Q78- and Q78 S2116A mutant-HTT (E) and Q23- and Q78 S2116A mutant-HTT (F) were assigned based on the equation described in the Materials and Methods and depicted as heat map. (G) Bar graph showing the number of unique cross-links between N-HEAT and C-HEAT domains from Q23- and Q78-HTT including corresponding S2116 alanine mutants.

To elucidate the effect of S2116A mutation on the global structure of HTT, we probed intramolecular interactions within Q23- and Q78-HTT and corresponding S2116A mutants by XL-MS using DSS as a cross linking reagent. This is a suitable technique for the characterization of the architecture of proteins especially since XL-MS can provide structural information within regions unresolvable through cryo-EM due to current technical limitations (Leitner et al., 2014a). The cross-linked peptides from all samples were identified by the xQuest search engine (Table S5) and the overall numbers were comparable (Table S6). Since HTT consists of three domains (Guo et al., 2018), we grouped the cross-linked peptide pairs into the three groups and analyzed the inter-domain (N-B, N-C, B-C) and intra-domain (N-N, C-C, B-B) changes in XL-MS analysis (Figure S15A). To quantify the differences, we assigned a fold enrichment score for each of the 6 types of interactions by considering the magnitude and direction o change of the cross-linked peptides (equation is presented in Materials and Methods). Consistent with previous XL-MS studies of HTT protein (Vijayvargia et al., 2016), the Bridge domain corresponding to the uncrosslinked domain (UCD) still showed unusually few cross-links (Figure S15B). Figure 7D shows a decrease in inter-domain interactions between the N-HEAT domain and the C-HEAT domain in Q78-HTT compared to those in Q23-HTT (−0.8 fold change), indicating an increase in distance between the N-HEAT and C-HEAT domains in Q78-HTT. This is surprisingly consistent with the cryo-EM results (Figure 4C) that also show a structural change resulting in increased separation of the N-HEAT and C-HEAT domains in Q78-HTT. In addition, the increased Q78-HTT N-HEAT intra-domain crosslinking (Figure 7D, 2.6 fold change) was mostly restricted to a subdomain of the N-HEAT domain that we call the NTD-III (previously called CTD-I) region (Figure S16) (Vijayvargia et al., 2016). This region is the pivot point linking the N-HEAT and Bridge domains (blue circles in Figure 5). These results are consistent with the idea that the Bridge domain serves as the pivot joint upon the polyQ tract expansion (Figure 5B). Thus, our XL-MS data supports and further specifies the structural changes between Q23- and Q78-HTT.

Most importantly, our XL-MS results revealed that the S2116A mutation specifically shifts the structural changes exhibited by Q78-HTT back towards Q23-HTT. S2116A mutation of Q78-HTT dramatically increased the inter-domain interaction between the N- and C-HEAT domains (Figure 7E), whereas the same mutation in Q23-HTT showed no or minimal impact (Figure S15B), indicating its mutant HTT specificity. When comparing Q23- and Q78 S2116A-HTT, we see that the C-HEAT—Bridge and N-HEAT—C-HEAT domain interactions of Q78 S2116A-HTT have shifted towards the pattern exhibited by Q23-HTT (Figure 7F). These results strongly suggest that S2116A mutation in the C-HEAT domain of mutant HTT may reinstate the intramolecular proximity between N-HEAT and C-HEAT domains toward normal HTT (Figure 7G) with a concomitant reduction of the functional consequences of the elongated polyQ tract on mutant HTT, implying the importance of the dynamic movement of the domains on HTT interaction.

## DISCUSSION

The *HTT* CAG expansion, encoding polyQ in HTT is a direct cause of HD. However, how the polyQ expansion in HTT affects its biological activity is obscure. Here, we propose a working model that the polyQ expansion alters the global structure of HTT and phosphorylation which is related to PRC2 stimulating activity of mutant HTT. Comparative structural analysis on our Q23/Q78 structures together with HTT_HAP40_ revealed several key insights about HTT structure and differences induced by polyQ expansion and protein binding.

First, superimposing N-HEAT domains from HTT_HAP40_, Q23-HTT and Q78-HTT clearly manifests a large conformational change among the domains with the N- and C-terminus of the Bridge domain acting as critical pivot joints (Figure 5A). In comparison to HTT_HAP40_, Q23-HTT exhibited an extended structure with the substantial conformational change in the CHEAT domain relative to the N-HEAT domain. Cryo-EM analysis of Q78-HTT showed a similar conformational change in the C-HEAT domain leading to an even more extended structure as compared to Q23-HTT, which is caused by polyQ expansion. Our data suggest a structural mechanism of HTT in response to HAP40 binding and polyQ length expansion (Figure 5B), which heavily involve modular movements of the C-HEAT domain relative to the N-HEAT domain. The C-HEAT domain moves outward from the N-NEAT domain in response to polyQ expansion, and moves inward to the N-HEAT domain in response to HAP40 binding. While large conformational changes occurs among the domains, detectable changes in structure are limited to the inter-domain movement, as shown by HDX-MS analysis. Hence, HTT maintains a modular structure while exhibiting dynamic movements in response to protein binding and polyQ expansion. Our comparative structural analysis of HTT proteins in different states strongly suggests that the conformational change of the C-HEAT domain relative to the N-HEAT domain may play critical roles in HTT function, and the polyQ expansion may regulate this conformational change leading to modulating HTT function.

Using a series of HTT proteins with different polyQ lengths, we showed that polyQ expansion affects the phosphorylation level of specific sites across the entire protein and many of these sites are located in disordered and/or surfaced-exposed regions, indicating that the phosphorylation sites are largely solvent accessible and able to interact with the polyQ tract and other potential binding partners. These data indicate that the polyQ expansion not only modulates the orientation of the modular structure of HTT, but also affects the biochemical characteristic of the disordered regions in HTT, including PTMs. Among many PTM sites, S2116 phosphorylation highlights how the interaction between the N- and C-HEAT domains can act as a critical feature for HTT function, both in the context of normal biological function and understanding the HD mechanism.

Considering that the polyQ expansion induces global structural changes by altering the conformation of the domains, this may lead to changes in PTM patterns that regulate HTT function by providing a binding platform for other proteins. Alternatively, PTMs may be directly regulated by the polyQ expansion, affecting the conformation of HTT and modulating HTT function. Based on our observations of S2116 and its influential role in HTT structure and function, we propose a complex, bidirectional relationship: that polyQ expansion may modulate the equilibrium between the domain movements and PTM patterns, which in turn further stabilizes and perpetuates the altered equilibrium of the domain movements. This dynamic configuration of HTT may be supportive by an elastic behavior of HEAT-repeat molecules suggested by molecular dynamic simulations of PR65A (Grinthal et al., 2010). A single phosphorylation site such as S2116 may act as a key regulator of cross-talk among multiple PTMs, which enact pleiotropic effects onto the protein, such as potentially reversing the numerous structural and functional changes brought about by polyQ expansion. To further evaluate the role of the phosphorylation at S2116, the cellular consequences of its phosphomutant need to be more intensively assessed, including identification of HTT phosphorylation status in a certain cell type. It should be noted that S2116 is located at the region interacting with HAP40 and phosphorylation of S2116 would affect the interaction between HTT and HAP40 (Figure S17), implicating the importance of S2116 for HTT function. The extent of HTT phosphorylation appears to be largely dependent on cell type and conditions, for example, S421, S1201 and S1181 sites were reported to be extensively phosphorylated in normal compared to mutant HTT (Anne et al., 2007; Warby et al., 2005). Our HTT phosphorylation PRM assays will provide a comprehensive evaluation of the phosphorylation levels of endogenous HTT with our HTT phospho-antibodies. It will be also critical to determine whether there are more cellular interactors that can be affected by the phosphorylation at S2116 in a polyQ length dependent manner. A recent study explored the role of several HTT PTM sites on mouse primary neuron toxicity caused by the polyQ expansion using nuclear condensation and mitochondria viability assays, suggesting no significant effect of S2116A, while some sites, such as S116 and S2652, inhibited the toxicity significantly (Arbez et al., 2017). However, this may indicate that the cellular consequence of phosphorylation depends on the specific target and cell status/type.

With the integration of structural, functional, and PTM characterization, we have built a proof-of-concept that polyQ expansion can exhibit structural consequences on HTT that can affect its functional activity in a modulatory fashion. From the various attempts to characterize HTT’s biological function, it is evident that HTT does not have a single, identifiable function or a well-defined set of binding partners, but rather participates in diverse cellular processes consistent with other HEAT-repeat proteins (Kobe et al., 1999). In fact, HEAT repeat protein PR65A can serve as a potential conceptual model for HTT: PR65A binds to two separate components, regulatory and catalytic, to build serine-threonine protein phosphatase 2A (PP2A) complex, and the interchangeable binding with the various regulatory domains influences PP2A enzymatic activity, allowing its various functions (Shi, 2009). Moreover, PR65A phosphorylation is suggested as an *in vivo* mechanism for regulating PP2A complex signaling (Kotlo et al., 2014). Similarly, HTT, through PTMs and/or conformational changes, may present slightly modified binding surfaces to other proteins, resulting in a dynamic proteomic milieu for HTT. HD therapeutics may require a more nuanced approach to ameliorating the toxicity of the mutation based on understanding various HTT functions, since the efficacy of gene therapy has not yet been confirmed and disease-modifying interventions may significantly precede the pre-symptomatic phase (Caron et al., 2018).

Overall the work presents a structural and molecular basis of the conformational flexibility of HTT mediated by polyQ expansion and phosphorylation and provides a lead for new strategies to understand the HD mechanism.

## MATERIALS AND METHODS

### Human FLAG-huntingtin cDNA clones

The information about FLAG-HTT cDNA cloned into pFASTBAC1 vector (Invitrogen) was mentioned in the previous paper (Vijayvargia et al., 2016). Additionally, for making eGFP fusion HTT clones, the Kas1 restriction site which is present in the 5’ end of HTT cDNA was used. The eGFP cDNA fragment was cloned into Kas1 site with In-Fusion Cloning Kit (Clontech). The eGFP-HTT construct contains a GSGS linker.

### Purification of HTT and eGFP fusion HTT

The purification of FLAG-tag HTT has been described (Vijayvargia et al., 2016). For purification of HTT, Baculovirus protein expression system (Invitrogen) was used. The Sf9 cell lysate was obtained by centrifugation at 39,000 x g, at 4 °C for 2 h and after freeze and thaw method in Buffer A (100 mM NaCl, 50 mM Tris-HCl pH 8.0, protease inhibitor cocktail (Roche), 5% Glycerol). The supernatant was incubated with M2 anti-FLAG beads (Sigma-Aldrich) for 2 h at 4 °C. After washing three times with Buffer A, bound proteins were eluted with Buffer B (100 mM NaCl, 50 mM Tris-HCl pH 8.0, 5% glycerol) containing 0.4 mg/ml Flag peptide. Eluted proteins were incubated in Buffer C containing150 mM NaCl, 50 mM Tris-HCl pH 8.0, 5% glycerol, 20 mM imidazole, 1 mM 1,4-dithiothreitol (DTT) and 0.5 mM EDTA with TEV protease for cleaving the N-terminal FLAG and 6X histidine tag (16 h, 4 °C). Tag cleaved HTT was further purified by Ni-NTA Agarose beads (QIAGEN) and Superdex 200 26/60 column (GE Healthcare) in running buffer (150 mM NaCl, 20 mM HEPES-HCl pH 7.5). To maintain the monomeric state of HTT, purified HTT was crosslinked with 0.5 mM DSS crosslinker for 30 min at room temperature. The monomeric fraction of HTT was separated by 10-30% sucrose density gradient fractionation. Crosslinked monomeric HTT was concentrated with Amicon Ultra 100-kDa filters (Milipore). Q78-HTT and GFP-Q23-HTT were purified by same procedure.

### Cryo-EM grid conditioning and image collection

For cryo-EM, the monomeric HTT (3.5 μl of 0.08 mg/ml) was mixed with final 0.05% of octyl glucoside and loaded on R2/1 300 mesh (Quantifoil) grids covered with graphene oxide (Bokori-Brown et al., 2016). The grids were incubated with HTT sample for 30 seconds, then vitrified by plunge freezing in liquid ethane using a Vitrobot II (FEI) with 8~10 seconds blotting time. The temperature and humidity of the vitrification chamber were maintained at 15 °C and 100% respectively. The grids were transferred to liquid nitrogen and loaded into an autoloader of Titan Krios microscope with a Gatan K2 Summit direct detector. Multiple data acquisitions were carried out for this study (Table 1). The images of Q23-, Q78-, and GFP-Q23-HTT were collected at the Diamond Light Source in the UK with a Volta phase plate (VPP). The detailed imaging conditions and parameters are summarized in Supplementary Table S1.

### Image processing and analysis

Movie frame stacks were imported into Scipion (de la Rosa-Trevin et al., 2016) to set a platform for accessing various program suites for further image processing. MotionCor2 (Zheng et al., 2017) was used for beam-induced motion correction and dose weighted and un-dose weighted micrographs were generated. The CTF estimation was done by using CTFFIND4 (Rohou and Grigorieff, 2015) and GCTF (Zhang, 2016), an option for detecting additional phase shift applied on VPP images. Good images were selected based on estimated CTF resolution (< 8 Å), defocus range (for VPP images, between −0.2 to 1.0 μm), and the thickness of graphene oxide. All micrographs were thoroughly inspected and the particle picking was carried out fully manually. The particles were extracted from dose weighted micrographs with a 250 x 250 pixel box size and the contrast was inverted for further image processing. The particle sets were cleaned by 4 to 8 rounds of reference free RELION 2D classification, and particles in good 2D classes were subjected to 3D initial model building using RELION 3D initial model. The initial maps (~ 20 Å resolution) were used as a starting model for refinement of the structure by using 3D classification and 3D auto refinement with selected particle sets in RELION (Figure S2, 4, and 6).

UCSF-Chimera (Pettersen et al., 2004) and Pymol (Alexander et al., 2011) were mainly used for further image analysis and validation. The estimated resolution was calculated from the EMDB FSC validation server (https://www.ebi.ac.uk/pdbe/emdb/validation/fsc/) using pairs of unfiltered half maps (Table S1).

### Integrative modeling of Q23 -HTT

The Q23-HTT was modeled at atomistic resolution using an integrative modeling strategy which consisted in (i) splitting the HTTHAP40 structure into three rigid subunits, (ii) rigidly fitting the subunits into the density map with a multi-body constrained molecular docking strategy and (iii) flexibly refining the atomistic models inside the density maps. Details of the distinct modeling steps are described in the following sections.

#### 1. Splitting HTT_HAP40_ structure into rigid subunits

The HTTHAP40 was split into three major subunits consisting in a N-HEAT domain (residue number 91 to 1684), bridge domain (residue number 1685 to 2091) and C-terminal HEAT domain (residue number 2092 to 3098). The missing residues corresponding to flexible regions of the HTT_HAP40_ were not directly included in the current modeling procedure.

#### 2. Rigid docking of HTT subunits inside cryo-EM maps

First, the N-HEAT domain subunit density map was rigidly fit with SITUS (Wriggers et al., 1999) inside the Q23-HTT cryo-EM density map. As a first step, we initially used the widely available SITUS package (Wriggers et al., 1999). Due to the fact the current version of SITUS could not simultaneously fit the three subunits, the fitting protocol was initiated by fitting the N-HEAT domain, which was the largest of the three partial models, into the density map using the SITUS module *colores*.

Later, the docking problem consisting in simultaneously fitting the remaining bridge and C-HEAT domains was modeled as a constrained optimization. The aim of this approach was to optimally fit the bridge and C-HEAT domains into the density map whilst avoiding steric clashes with the already SITUS-fitted N-HEAT domain and keeping their native intersubunit connectivity.

To perform such a task, the mViE (memetic viability evolution) algorithm was coupled to our in-house docking protocol *pow^er^* (parallel optimization workbench to enhance resolution). The combination of *pow^er^* and mViE to solve integrative modeling problems has been thoroughly described in (Tamo et al., 2017) and thus only a brief explanation is given here. *Pow^er^* is an integrative modeling platform that was designed to virtually solve any type of optimization problems (Degiacomi and Dal Peraro, 2013). It has been successfully applied to solve a multitude of protein assembly problems (Tamo et al., 2015) by using state-of-the-art heuristic optimization algorithms to efficiently sample the large search space associated with molecular docking (Degiacomi and Dal Peraro, 2013).

Amongst the list of optimizers featured in *pow^er^*, mViE is a recently implemented constraint optimization algorithm that evolves a population based on (1+1)-covariance matrix adaptation evolution strategy (CMA-ES) for local search of the search space, which can recombine using differential evolution for global search (Floreano, 2015). To optimize the balance between local and global search, mViE uses a dedicated task scheduler. Importantly, mViE was shown to outperform several state-of-the-art constrained optimizers on a standard constrained optimization benchmark set (Floreano, 2015). The combination of *pow^er^* with mViE was recently shown to effectively solve the problem of fitting macromolecular assemblies inside density maps (Tamo et al., 2017), and thus was used in this work to model the Q23-HTT structure using structural information from the experimental cryo-EM density map.

The process resulting in the optimal fit of HTT subunits into the map was encoded as a contrained optimization problem. In this context, the search space sampled by power-mViE consisted in the orientation and position of each HTT subunit which were encoded as [*x_Nterm_, y_Nterm_, z_Nterm_, α_Nterm_, β_Nterm_, γ_Nterm_, x_bridge_, y_bridge_, z_bridge_, α_bridge_, β_bridge_, γ_bridge_*] where *x, y*, and *z* corresponded to three translation sampled by the N-HEAT domain and bridge respectively; and *α, β and γ* the Eulerian angles defining the spatial orientation of respective domains. These spatial parameters were simultaneously sampled in order to solve the problem formulated as:

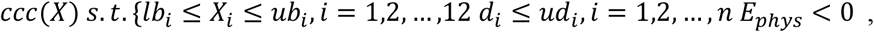

Where 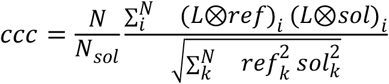 corresponded to the CCC quantitatively estimating the fit between the assembled HTT subunits and the experimental cryo-EM map. In this case, L corresponds to a 3×3×3 Laplacian filter kernel used to convolute the input density maps with the aim to increase the precision of the density map fitting. *N* is the number of overlapping voxels between the density maps, *N_sol_* the total number of voxel of the candidate solution density map, (*L* ⊗ *ref*)_*i*_ is the *i*^th^ voxel of the reference density map and (*L* ⊗ *sol*)_*i*_ its counterpart from the candidate monomer assembly.

The CCC was maximized upon the satisfaction of pre-defined inequality constraints. The inequality constraints were defined by *lb_i_* and *ub_i_*, which were the lower and upper lower bound defining the search space of *X_i_*; *ud_i_* which corresponded the upper Euclidian distances on Cα atoms respectively defining the spatial connectivity between residue 1713 – 1729, residue 2062 – 2092, and between two distances extracted from XL-MS experiments which were residue 943 - 1973 and residue 882 - 1882 to correctly orient the Bridge domain relative to the N-HEAT domain. These distance constraints were intentionally chosen to be < 45 Å as they were used to coarsely guide the assembly process, rather than provide accurate positions; 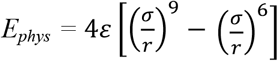 is a 9-6 Lennard-Jones coarse energy potential (Alemani et al., 2010) used here to prevent steric clashes, where r consists in the pairwise distance between the Cα interfacial atoms of the subunits within a distance of 12 Å, *σ* = 4.7 Å and *ε* = 1 kcal/mol. In order to ensure a thorough sampling of the search space, a total budget of 1.8 million function evaluation was allocated to the power-mViE optimization protocol. The best model ranked by constraint satisfaction and CCC value was extracted and further refined.

#### 3. Flexible refinement of the docked subunits inside cryo-EM maps with MDFF

The fit of the best-scoring structures inside the density map was refined using the Molecular Dynamics Flexible Fitting (MDFF) protocol (Trabuco et al., 2009). The simulations for Q23-HTT were performed using NAMD (Phillips et al., 2005) *in vacuo* using the CHARMM forcefield (MacKerell et al., 2000) with a dielectric constant of 80. The SHAKE algorithm was applied to restrain all bonds and secondary structure restraints were used to prevent the structures from unraveling during the refinement process.

### Integrative modeling of Q78-HTT

The knowledge acquired from fitting the Q23-HTT domains positions enabled to detect the approximate location of N-, C-term and bridge domain inside Q78-HTT density map. Due to the low-resolution of Q78-HTT cryo-EM map at the location of Q78-HTT bridge domain, the bridge structure was omitted from the final Q78-HTT model.

The rigid fit of both Q78-HTT N- and C-terminal domains was performed in successive steps. First, N-HEAT domain of the Q23-, and Q78-HTT cryo-EM map were aligned. Owing to this alignment, the Q23-HTT N-terminal structure of Q23 could be readily transferred from Q23-HTT EM map into Q78-HTT map using the UCSF-Chimera Fit in map command. The Q78-HTT C-HEAT was then manually inserted into its corresponding C-terminal map region, and locally fitted with the UCSF-Chimera “Fit in map” command. This initial rigid fit of N-HEAT and C-HEAT was used as seed for a more refined fit into cryo-EM map using MDFF protocol (Trabuco et al., 2009). The simulations for Q78-HTT were performed using NAMD (Phillips et al., 2005) *in vacuo* with the same parameters used in Q23-HTT simulation. The simulation lasted 600 ns until Cα-atom converged, and the CCC value of MDFF-refined Q78-HTT structure against the cryo-EM map was 0.84 (Figure S5B).

### Data availability

EMDB ID for Q23-HTT, GFP-Q23-HTT, and Q78-HTT are EMD-4937, EMD-4944, and EMD-4946, respectively. PDB ID for Q23-HTT MDFF model: 6RMH. roteomeXchange/PRIDE. ID PXD013907 for XL-MS.

### Hydrogen-deuterium exchange mass spectrometry

The labeling reaction was started by mixing 3 μL of each protein batch (Q23-, or Q78-HTT) with 23 μL of deuterated buffer. For kinetics, deuteration time was 10 min, 45 min, 90 min, and 180 min (all incubations were done at room temperature). Each time point of the deuterium-labeling reaction was stopped by decreasing the pH to ~ 2.3 and temperature to ~ 4 °C by means of adding 30 μL of a solution containing 250 mM TCEP, 3 M Urea and 200 mM phosphate buffer pH 2.3.

Each labeled and quenched sample was analyzed in a semi-automated HDX-MS system (Biomotif AB, Stockholm, Sweden) in which manually injected samples were automatically digested, cleaned and separated at 2 °C. Deuterated samples were digested using a Poroszyme immobilized pepsin column (Applied Biosystems 2.1 mm x 30 mm) by a 1 min at 70 μl/min flow protocol, followed by an on-line desalting step using a 2 mm I.D x 10 mm length C-18 precolumn (ACE HPLC Columns, Aberdeen, UK) using 0.05% TFA at 250 μl/min for 3 min. Peptic peptides were then separated by the following gradient (Buffer A consisted of 0.3 % formic acid and 5 % acetonitrile and buffer B consisted of 0.3 % formic acid, 95 % acetonitrile and 5 % water): 5 % B buffer at 0 min, 20 % B buffer at 7 min, 35 % B at 15 min, 90 % B in 20 min and 5 % B at 25 min using a Chromolith FastGradient RP-C-18, 50 mm length and 2 mm diameter analytical column (Merck) operated at 120 μl/min. An Orbitrap XL mass spectrometer (Thermo Fisher Scientific) operated at 60,000 resolution at m/z 400 was used for analysis. For peptide identification, several LC MS/MS runs were performed and searched against an in-silico database consisting of the Q23-, or Q78-HTT sequence. All obtained HDX-MS data were analyzed by the HDExaminer software (Sierra Analytics, USA).

### Small-angle X-ray scattering

The data were collected at the European Synchrotron Radiation Facility (ESRF), beamline BM29 (Pernot et al., 2013). The samples were concentrated to 8.8 mg/ml and were loaded onto a Superose 6 10/300 GL column (GE Healthcare) equilibrated with 20 mM HEPES, pH 7.5, 150 mM NaCl, 10% glycerol and 2 mM TCEP. The flow rate was set to 0.5 ml/min and the temperature to 18 °C. As they were eluted from the column, the samples were illuminated with X-rays of wavelength 0.99 Å. A PILATUS 1M detector was used at a distance to the sample of 2.86 m covering a momentum transfer of 0.025 < q > 5 nm-1 (q = 4πsin (θ)/λ, where θ is the total scattering angle). During the 50 min elution, 3000 scattering measurements were taken with 1s time-frames. SAXS data were analyzed using the ATSAS package and ScÅtter (Franke et al., 2017; JA, 2011). For each sample, using CHROMIXS (chromatography in-line X-ray scattering), an elution profile was generated with the integrated intensities plotted versus the recorded frame number. In CHROMIXS, ~50 buffer frames were averaged and used to (i) subtract the buffer average from each frame of the sample peak selected and (ii) calculate the corresponding Radius of Gyration (Rg). Only measurements with equal Rg values were included in the final sample peak region chosen. The subtracted sample peak region was averaged to generate the final scattering curve used for subsequent analysis. The scattering curves were initially viewed in PRIMUS (P. V. Konarev, 2003) where the Rg was obtained from the slope of the Guinier plot (Guinier, 1939) within the region defined by qmin < q < qmax where qmax < 1.3/Rg and qmin is the lowest angle data point included by the program. The P(r) function, the distribution of the intra-atomic distances (r) in the particle, was generated using the indirect transform program GNOM (P. V. Konarev, 2003). The maximum distance (Dmax) was selected by letting the P(r) curve decay smoothly to zero. DATPOROD, DATOW, and DATVC (H. Fischer, 2010; M. V. Petoukhov, 2007; Rambo and Tainer, 2013) were used to estimate the Porod Volume (Vp) and the concentration-independent estimate of the MW for the samples.

### Mass spectrometric identification of phosphorylation sites

For the determination of phosphorylation sites, recombinant full-length human HTT proteins with polyglutamine tract length of 23, 46 and 78 were purified in the presence of complete protease and phosphatase inhibitor cocktails (Roche Applied Science, Branford, CT) to retain phosphorylation. 10-20 μg of purified proteins were separated by SDS-PAGE and stained with mass spectrometry compatible GelCode Blue stain reagent (Thermo Scientific, Carlsbad, CA). HTT bands were excised from gel and processed for mass spectrometry. The samples were reduced with 1 mM DTT for 30 min at 60°C and then alkylated with 5 mM iodoacetamide for 15 min in the dark at room temperature. Gel pieces were then subjected to a modified in-gel trypsin digestion procedure (Shevchenko et al., 1996). LC-MS/MS analysis of the digests was carried out on an LTQ-Orbitrap mass spectrometer (ThermoFinnigan, San Jose, CA). The eluted peptides were detected, isolated, and fragmented to produce a tandem mass spectrum of specific fragment ions for each peptide. Peptide sequences (and hence protein identity) were determined by matching protein or translated nucleotide databases with the acquired fragmentation pattern by the software program TurboSEQUEST v.27 (ThermoFinnigan, San Jose, CA) (Eng et al., 1994). The modification of 79.9663 mass units to serine, threonine, and tyrosine was included in the database searches to determine phosphopeptides. Each phosphopeptide that was identified by the SEQUEST program was also manually inspected to ensure confidence. Phosphopeptides obtained were manually aligned to the HTT sequence to generate the coverage map. Amino acid count in peptides was plotted to determine the sequence coverage.

### Antibodies

HTT phosphorylation site specific affinity purified rabbit polyclonal antibodies against the 16 sites identified in this study were generated by New England Peptide (Gardner, MA). In short, KLH-coupled peptides bearing the specific phosphorylated residue were injected into 2 rabbits each. Booster doses were given on day 14 and 28. Production bleeds were checked for antibody titer by ELISA using purified phosphopeptides. High-titer bleeds (40-50 mL) were subjected to a double affinity purification. The first step involved negative selection wherein affinity columns with non-phosphorylated peptide backbones were used. The antibodies against the peptide backbone are retained in the column while those against the phosphorylated residue are in the flow through. The flow through is then subjected to a second cycle of affinity purification (positive selection) wherein phosphorylated peptides are used to specifically interact with phosphorylation site specific antibodies. The bound antibodies are eluted by lowering the pH. The specificity of phosphorylation site specific antibodies was validated by a dot-blot assay. Briefly, 100 ng of phosphorylated and non-phosphorylated peptides were spotted onto a nitrocellulose membrane and allowed to dry. The membrane was blocked with 5% BSA/TBST and probed overnight with 0.25 μg/mL of different phosphorylation site specific antibodies in 5% BSA/TBST at 4°C. After washing, the blots were probed with anti-Rabbit HRP secondary antibodies and developed using the ECL system. Anti-HTT mouse monoclonal antibody mAb2166 was purchased from Millipore (EMD Millipore, Darmstadt, Germany).

### Immunoblot analysis

50-100 ng of purified protein was run on a 6% or 10% Bis-Tris gel and transferred onto nitrocellulose membranes. All antibodies were blocked with 5% milk/TBST. However, phosphorylation site specific antibodies were used at 0.25 μg/mL in 5% BSA/TBST and incubated overnight at 4°C for probing. Anti-HTT antibodies were used at dilutions of 1:2000 (mAb2166). The data obtained by western blotting were quantified by subjecting the scanned images to densitometry using Image J software (NIH).

### In gel tryptic digestion of endogenous HTT from human lymphoblastoid cell lines (LCL)

Gel slices were cut into 1×1 mm pieces and placed in 1.5 mL eppendorf tubes with 1 mL of water for 30 min. The water was removed and 50 μL of 250 mM ammonium bicarbonate was added. For reduction 10 μL of a 45 mM solution of DTT was added and the samples were incubated at 50°C for 30 min. The samples were cooled to room temperature and then for alkylation 10 μL of a 100 mM iodoacetamide solution was added and allowed to react for 30 min. The gel slices were washed 2 times with 1 mL water aliquots. The water was removed and 1 mL of 50:50 (50 mM ammonium bicarbonate: acetonitrile) was placed in each tube and samples were incubated at room temperature for 1 h. The solution was then removed and 200 μL of acetonitrile was added to each tube at which point the gels slices turned opaque white. The acetonitrile was removed and gel slices were further dried in a Speed Vac. Gel slices were rehydrated in 70 μL of 2 ng/μL trypsin (Sigma) in 0.01% ProteaseMAX Surfactant (Promega): 50 mM ammonium bicarbonate. Additional bicarbonate buffer was added to ensure complete submersion of the gel slices. Samples were incubated at 37 °C for 21 h. The supernatant of each sample was then removed and placed in a separate 1.5 mL eppendorf tube. Gel slices were further dehydrated with 100 μL of 80:20 (acetonitrile:1% formic acid). The extract was combined with the supernatants of each sample. The samples were then dried down in a vacuum centrifuge. Samples were dissolved in 30 μL of 5% acetonitrile in 0.1% trifluoroacetic acid prior to LC-MS/MS analysis.

### HTT phosphopeptides quantification using parallel reaction monitoring (PRM) assay

A 3.0 μL aliquot was directly injected onto a custom packed 2 cm x 100 μm C_18_ Magic 5 μm particle trap column. Peptides were then eluted and sprayed from a custom packed emitter (75 μm x 25 cm C_18_ Magic 3 μm particle) with a linear gradient from 95% solvent A (0.1% formic acid in water) to 35% solvent B (0.1% formic acid in Acetonitrile) in 60 min at a flow rate of 300 nL per minute on a Waters Nano Acquity UPLC system. Ions were introduced by positive electrospray ionization via liquid junction into a Q Exactive hybrid mass spectrometer (Thermo Scientific). Data-dependent acquisition (DDA) experiments for library generation were performed according to an experiment where full MS scans from 300-1750 m/z were acquired at a resolution of 70,000 followed by 10 MS/MS scans acquired under HCD fragmentation at a resolution of 17,500 with an isolation width of 1.6 Da. For parallel reaction monitoring (PRM) a total of 13 m/z’s representing phosphorylated and control peptides observed in the DDA analyses were continuously monitored throughout the gradient using an isolation width of 2 Da followed by HCD fragmentation at a collision energy of 27% acquiring MS/MS spectra at a resolution of 17,500. A PRM analysis run in technical triplicates was performed on a cohort of 24 HD patient samples. Raw data files were processed with Proteome Discoverer (Thermo, version 1.4) prior to searching with Mascot Server (Matrix Science, version 2.5) against a human database which contained both variants of the HTT protein. Search parameters utilized were fully tryptic with 2 missed cleavages, parent mass tolerances of 10 ppm and fragment mass tolerances of 0.05 Da. A fixed modification of carbamidomethyl cysteine and variable modifications of acetyl (protein N-term), pyro glutamic for N-term glutamine, oxidation of methionine, and phosphorylation of Ser/Thr were considered. Quantitation of targeted peptides was accomplished through the use of the Skyline software (University of Washington, version 3.5). Spectral libraries used by Skyline were constructed from Proteome Discoverer search result MSF files generated from DDA runs. Top three fragment ion intensities were extracted from each of the 7 peptides and summed to determine the peptide abundance.

### Cell-free assay for PRC2 methyltransferase activity

Reconstituted *in vitro* PRC2 activity assays consisting of G5E4 DNA with 12 nucleosomes were performed essentially as previously described (Seong et al., 2010) using 20 nM of full-length HTT (Q23 and Q78 including corresponding alanine mutants) to modulate phosphorylation status with 10 nM of PRC2 complex.

### Immunoprecipitation

To determine the effect of HTT phosphorylation status on its interaction with PRC2 complex, 2 μg of HTT proteins (Q23- and Q78-HTT including corresponding S2116A mutants) were mixed with 2 μg of purified PRC2 complex or buffer and incubated together for 2 h at 4°C in 100 μL of 50 mM Tris-HCl pH 8.0, 150 mM NaCl, and 0.5 mM EDTA. 1-2 μg of anti-EZH2 antibody was added subsequently and further incubated overnight at 4°C. The antigen-antibody complex was pulled down by the addition of 50 μL of protein-G agarose slurry (Roche Applied Science, Branford, CT). Non-specific binding was removed by excessive washing of the beads (6 times) with wash buffer (50 mM Tris-HCl pH 8.0, 300 mM NaCl, 0.5 mM EDTA) and the complex was eluted by adding 50 μL of Laemmli sample buffer followed by boiling at 95°C for 5 min. The eluted samples were separated by SDS-PAGE and subjected to immunoblotting with anti-HTT antibody.

### Cross-linking Mass Spectrometric analysis of HTT

#### 1. Preparation of cross-linked HTT monomer

Cross-linking of full-length HTT was carried out as previously described (Vijayvargia et al., 2016). Briefly, HTT was cross-linked with 0.5 mM of DSS-H12/D12 (Creative Molecules) for 30 min at 25°C. The reaction was quenched by adding 20 mM of ammonium bicarbonate. Monomeric fraction of HTT was separated with 10-30% sucrose gradient by ultracentrifugation at 111,541 x g for 16 h at 4°C.

#### 2. Sample processing for MS analysis

Cross-linked and quenched samples were dried in a vacuum centrifuge and redissolved in 50 μL of 8 M urea. Cysteines were reduced by addition of tris(2-carboxyethyl)phosphine to a final concentration of 2.5 mM and incubation for 30 min at 37 °C with mild shaking. Free thiol groups were subsequently carbamidomethylated by addition of iodoacetamide to 5 mM and incubation at room temperature for 30 min in the dark. Alkylated samples were diluted to 5.5 M urea by addition of 150 mM ammonium bicarbonate and 500 ng of endoproteinase Lys-C (Wako) were added, followed by incubation for 3 h at 37 °C. The samples were then further diluted to 1 M urea by addition of 50 mM ammonium bicarbonate, 1 μg of trypsin (Promega) was added and samples were incubated overnight at 37 °C. Digestion was stopped by addition of formic acid to 2% (v/v) and samples were cleaned up using 50 mg Sep-Pak tC18 cartridges (Waters). Eluates were evaporated to dryness in a vacuum centrifuge and resuspended in 20 μL water/acetonitrile (70:30, v/v) for fractionation by size exclusion chromatography (SEC) (Leitner et al., 2012). SEC was performed on an Äkta micro system equipped with a Superdex Peptide PC 3.2/300 column (both GE Healthcare) and a mobile phase consisting of water/acetonitrile/trifluoroacetic acid (70:30:0.1, v/v/v), operating at a flow rate of 50 μL/min. Three 100 μL fractions corresponding to the elution window of cross-linked peptides were collected separately and evaporated to dryness.

#### 3. Liquid chromatography-tandem mass spectrometry (LC-MS/MS)

Dried SEC fractions were redissolved in 20 μL of water/acetonitrile/formic acid (95:5:0.1, v/v/v) and 4 μL were used for LC-MS/MS analysis. All injections were performed in duplicate. LC-MS/MS was performed using an Easy-nLC 1000 nanoflow HPLC system coupled to an Orbitrap Elite mass spectrometer equipped with a Nanoflex source (all ThermoFisher Scientific). Peptides were separated on an Acclaim PepMap RSLC C18 column (15 cm × 75 μm, 2 μm particle size, ThermoFisher Scientific) using mobile phases A (water/acetonitrile/formic acid, 95:5:0.15, v/v/v) and B (acetonitrile/water/formic acid, 95:5:0.15, v/v/v) with a gradient of 9 to 35%B in 90 min and a flow rate of 300 nL/min. Data acquisition was configured with an MS scan in the Orbitrap analyzer (120,000 resolution), fragmentation of selected precursors in the linear ion trap (isolation with of 2.0 m/z, normalized collision energy of 35%) and acquisition of 10 dependent MS/MS scans in the linear ion trap at normal resolution. Dynamic exclusion was activated with an exclusion duration of 30 s and a repeat count of 1.

#### 4. Identification of cross-linked peptides with xQuest

Cross-linked peptides were identified with xQuest, version 2.1.4 (Leitner et al., 2014b; Rinner et al., 2008). Searches were performed against a database consisting of the huntingtin Q23 wild-type sequence and 13 contaminants (tubulins, heat shock proteins and human keratins). Search settings included enzyme = trypsin, maximum number of missed cleavages = 2, mass tolerance = 15 ppm for MS data and 0.2/0.3 Da for MS/MS data. All candidate intra-protein cross-link identifications with a score ≥16 were further filtered with a narrower mass tolerance of ±5 ppm and a %TIC subscore ≥0.1. All remaining spectra were manually evaluated and only assignments with at least four bond cleavages overall or three consecutive bond cleavages per peptide were considered further. Because the likelihood of false positives for intra-protein crosslinks at this score threshold is very low (Walzthoeni et al., 2012), we expect a false discovery rate of <1% for these datasets. For all samples combined, only five cross-links on contaminants were identified in total. The identified cross-linked pairs were grouped based upon lysine residues in the primary HTT sequence and unique cross-links number was counted. The fold enrichment score for each type of interaction was calculated based on the equation below:

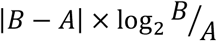

A: Number of identified cross-linked pairs of Q23- or Q78-HTT

B: Number of identified cross-linked pairs of Q78- or Q78 S2116A-HTT

## ACKNOWLEDGEMENTS

This work is partially supported by grants [NRF-2016K1A1A2912057 and NRF-2016R1A2B3006293 to J.S.] by the National Research Foundation (NRF) of Korea, and also supported by National Institutes of Health/National Institute of Neurological Disorders and Stroke [R01 NS079651 and R03 NS108028 to ISS]; CHDI foundation to I.S.S. T.J. and H.H. were supported by the Swedish Research Council (project grant 2016-03810). The group of R.A. was supported by European Research Council (ERCaG grant 670821 PROTEOMICS4D). The group of M.D.P. was supported by the Swiss National Science Foundation (grant number 200021_157217). This work was partially supported by the Intelligent Synthetic Biology Center(ISBC) of Global Frontier Project funded by the Ministry of Science and ICT(MSIT) (2011-003955). We are grateful to Dr. Ross Tomaino (Taplin mass spectrometry facility) and Dr. Jon Leszyk (UMASS medical school mass spectrometry facility) for excellent technical support. We also thank for the beam line scientist at the Beamline BM29 at European Synchrotron Radation Facility (ESRF). The computing resource for cryo-EM analysis was supported by Global Science experimental Data hub Center (GSDC), Korea Institute of Science and Technology Information (KISTI) [NRF-2018R1A6A7052113 to J.S.]. For data collection, we acknowledge the staffs at the UK national electron bio-imaging centre (eBIC), Diamond, funded by the Wellcome Trust, MRC and BBSRC and at the SciLifeLab, Swedish National cryo-EM Facility, funded by the Knut and Alice Wallenberg, Family Erling Persson and Kempe Foundations.

